# A new model for ancient DNA decay based on paleogenomic meta-analysis

**DOI:** 10.1101/109140

**Authors:** Logan Kistler, Roselyn Ware, Oliver Smith, Matthew Collins, Robin G. Allaby

## Abstract

The persistence of DNA over archaeological and paleontological timescales in diverse environments has led to revolutionary body of paleogenomic research, yet the dynamics of DNA degradation are still poorly understood. We analyzed 185 paleogenomic datasets and compared DNA survival with environmental variables and sample ages. We find cytosine deamination follows a conventional thermal age model, but we find no correlation between DNA fragmentation and sample age over the timespans analyzed, even when controlling for environmental variables. We propose a model for ancient DNA decay wherein fragmentation rapidly reaches a threshold, then subsequently slows. The observed loss of DNA over time is likely due to a bulk diffusion process, highlighting the importance of tissues and environments creating effectively closed systems for DNA preservation.

## Introduction

The genomic era of massively parallel DNA sequencing has driven a revolutionary body of research using ancient DNA-based genomics (1, 2). Paleogenomics has led to the re-writing of recent hominin evolutionary history (3), nuanced understandings of historical human movements and interactions around the globe (4, 5), breakthroughs in Quaternary paleontology (6–8), evolutionary ecology, the biology of extinct species (9), impacts of humans on ancient ecosystems and biodiversity (10), and the evolution and movements of domestic plants and animals (11–14). The successful probing of ancient epigenomes, microbiomes, and metagenomes further illustrates the flexibility and information value of ancient DNA-based research in the genomic age (15–17). In sum, time-series genomic datasets have proven extremely valuable in diverse research avenues.

In addition to the scale and sensitivity of analysis afforded by genomic methods in ancient DNA research, genomic datasets allow for a revised understanding of the patterns and expectations of DNA survival over millennia. This is beneficial in two key ways: i) Criteria of ancient DNA authenticity warrant updating for the genomic era, and formalized expectations of DNA degradation are necessary for this process; and ii) Better predictive models of DNA degradation may help researchers target specimens likely to yield high information value where destructive analysis is unavoidable. Generally, ancient DNA is expected to be highly fragmented (18) and to carry an abundance of characteristic misincorporations—deaminated cytosine residues appearing as C-to-T transitions in single-stranded fragment overhangs (19). Further, DNA fragmentation is biased by biomolecular context. For example, a short-range (∼10bp) periodicity observed in the distribution of fragment lengths is attributed to the period of a complete turn of the DNA double-helix around a histone (20), which is thought to offer some protection against breakage at histone-adjacent sites. Finally, base compositional biases have been regularly observed in DNA preservation, especially enrichment of GC-content in ancient DNA (21).

The relationships between these characteristic patterns of DNA degradation and the preservational environment and age of tissues are poorly understood. We carried out a meta-analysis of 185 ancient genomic datasets—dating from the Middle Pleistocene to the nineteenth century from 21 published studies (Figure 1; data sources cited fully in Supplemental Methods)—to test for relationships between sample age, environmental variables, and DNA diagenesis.

**Figure 1.**
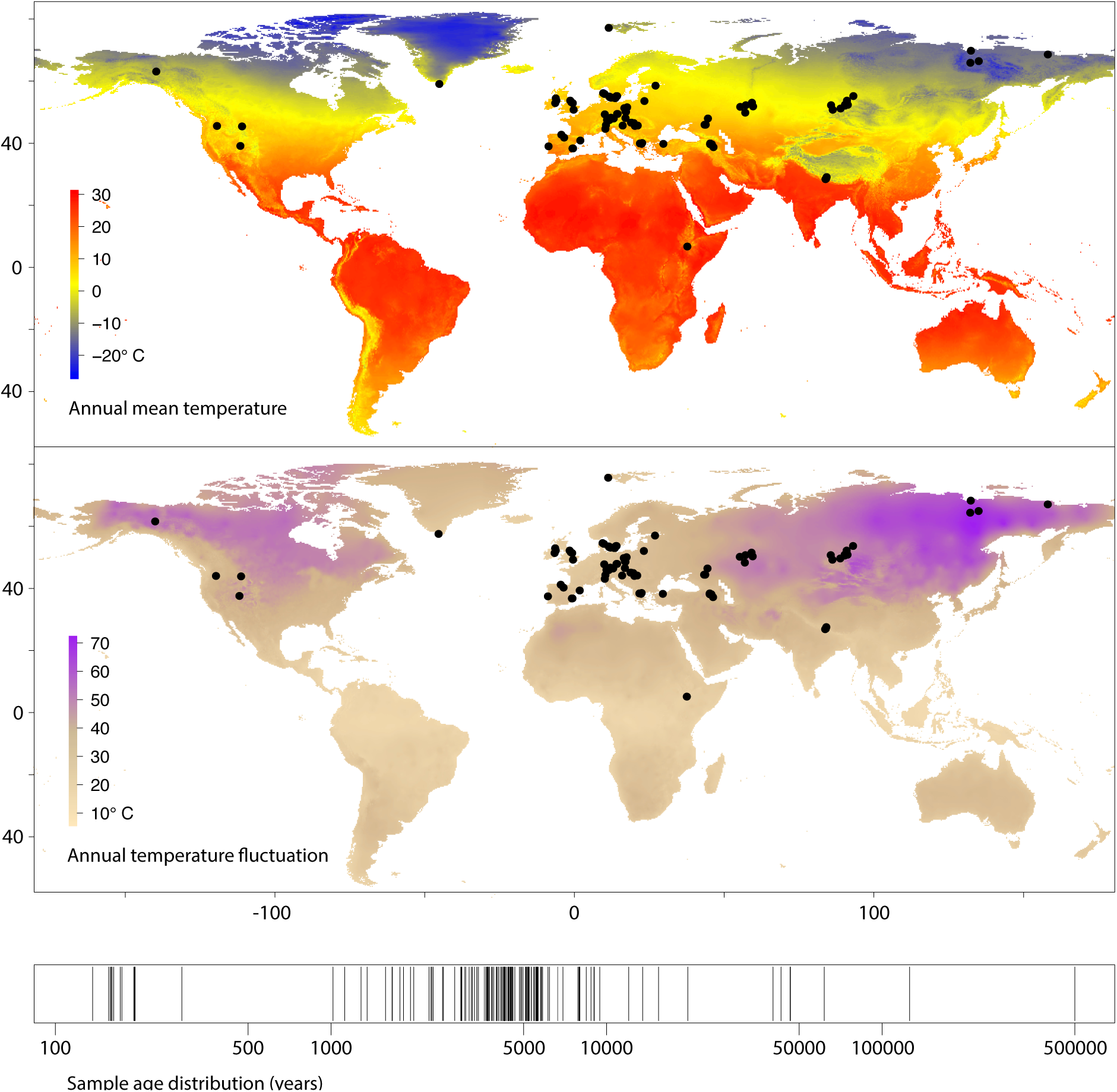
Locations of 185 samples (n=94 unique sites) used in paleogenomic meta-analysis, global variation in mean temperature and temperature fluctuation, and timeline of sample ages. Note the absence of sites with annual mean temperature >20°C, reflecting known preservation bias toward cooler climates (22).

We used mapDamage 2.0 (23) to quantify deamination, and we developed tests for assessing fragmentation, histone periodicity, and energetic biases in ancient genomic data (described in Supplemental Methods). We analyzed these damage statistics in relation to sample age, annual mean temperature, temperature fluctuation, and precipitation—treated as a proxy for humidity—using simple multivariate linear models (Supplemental Methods). Ultimately, we aimed to establish the key determinants of DNA survival, and the specific patterns of DNA breakdown expected under variable conditions.

## Results and Discussion

We found that cytosine deamination is strongly influenced by both sample age and site mean temperature (multiple r^2^ = 0.264; age *p* = 1.9 × 10^−9^; temperature *p* = 1.52 × 10^−5^, model *p* = 2.54 × 10^−10^; Figure 2). Previous studies have identified age as the key critical predictor of deamination (24), but our finding is in line with predictions of a time-dependent hydrolytic process where activation energy is achieved more frequently at higher ambient temperatures. A rate of deamination can be calculated for any sample with a known age and partial conversion of exposed cytosines (Supplemental Methods; Figure 3). The resulting rates vary widely, and show a strong correlation with temperature (r^2^ = 0.279; *p* = 1.23 × 10^−12^). In sum, deamination is a time-dependent process heavily modulated by temperature. When analyzing DNA fragmentation, however, we found that precipitation and thermal fluctuation were strong predictors (multiple r^2^ = 0.202; precipitation *p* = 0.0025; temperature fluctuation *p* = 6.18 × 10^−8^) but that age was not significantly correlated with the degree of fragmentation (*p* = 0.77), even when controlling for environmental conditions.

**Figure 2.**
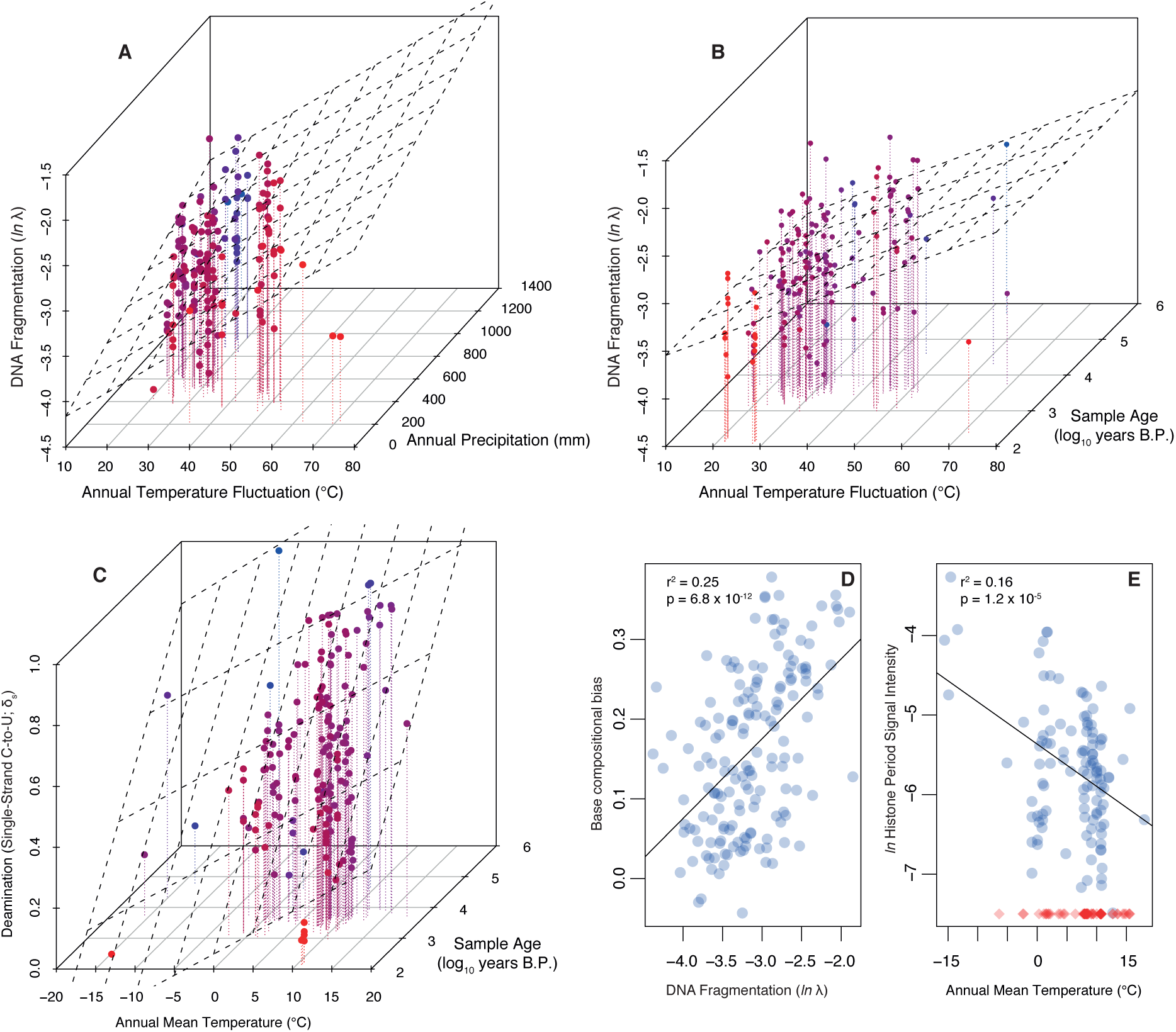
Relationships between DNA degradation parameters and environmental variables. **A)** DNA fragmentation is correlated with thermal fluctuation and precipitation. **B)** DNA fragmentation is correlated with thermal fluctuation, but is not influenced by sample age. **C)** Deamination is a thermal age parameter, strongly associated with both age and temperature. Coloring in A-C is used to enhance the z-axis variation: Red points are the nearest and blue are the most distant. **D)** DNA fragmentation is highly predictive of base compositional biases, with fragmented datasets depleted of motifs with low base-stacking energy. **E)** Histone periodicity in fragment length distribution is most pronounced in samples from cold environments. Blue circles represent samples where a histone periodicity estimate was possible (n=112; see Supplemental Methods for calibration against false positive results), red diamonds are samples where no periodicity was observed, visualized at -7.5 on the y-axis to reflect the observation of no detectable bias.

**Figure 3.**
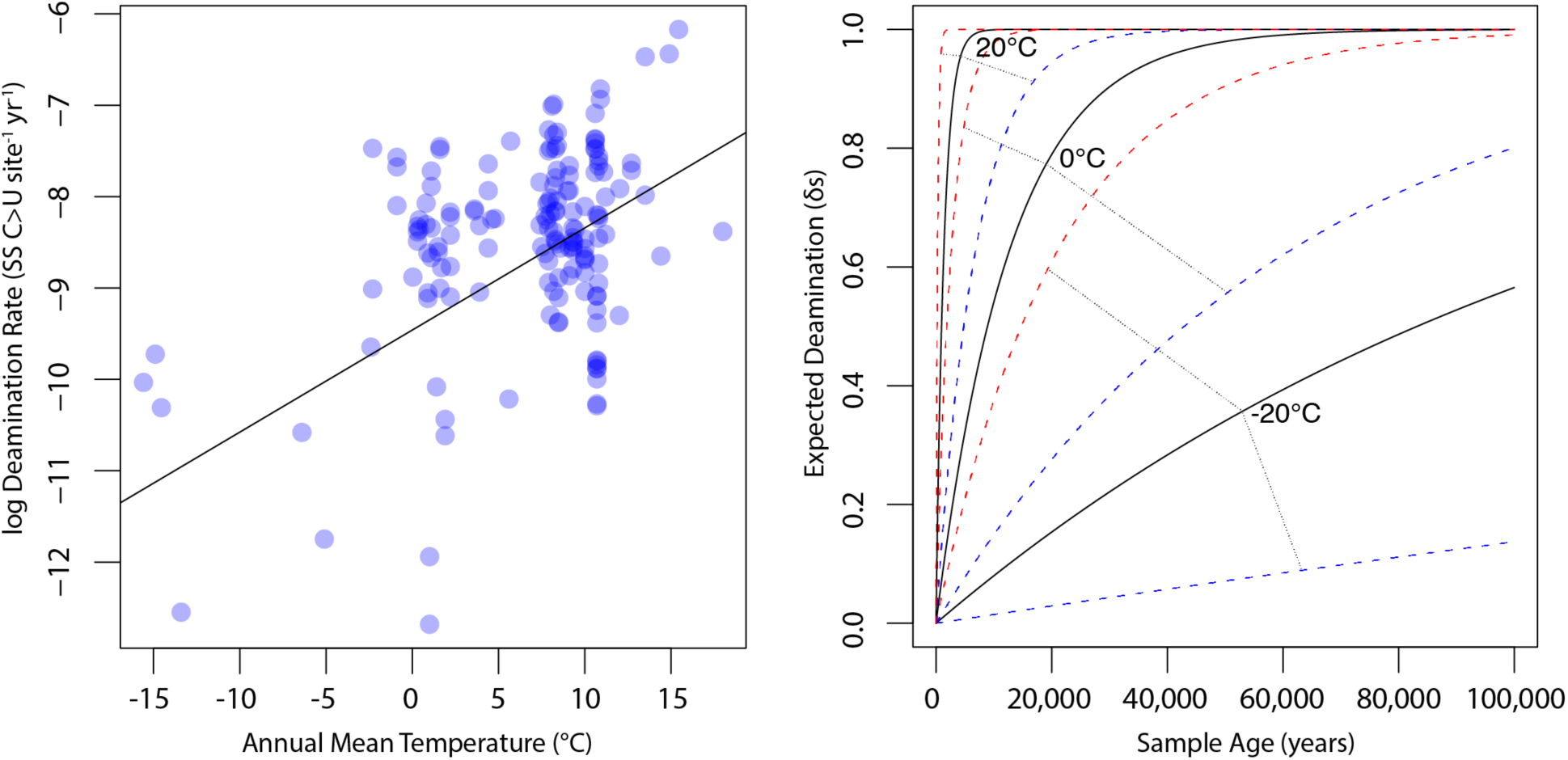
Expectations of deamination over time for variable temperatures. Left: Density-weighted linear regression of temperature and log deamination rate calculated using the formula 
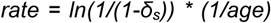
. Right: Using rate estimates from the weighted regression, we calculated the expected δ_s_ values over a 100,000 year timespan using the formula 
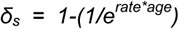
 re-arranged from the above, as well as δ_s_ values for rates estimated from 95% confidence intervals of the regression. We visualized the expected deamination levels for samples from -20°C, 0°C, and 20°C contexts (solid lines), along with upper (red) and lower (blue) confidence bounds. This predictive model is necessarily based primarily on mammalian bone tissue, and we expect refinements to these expectations based on sample type, for example, as more datasets become available.

At present, the ancient DNA literature lacks clear consensus concerning some of the fundamental predictors of DNA fragmentation: One recent study identified a strong age dependency in DNA recovery through qPCR analysis of a regionally controlled time series of bone samples (18). This result was interpreted as evidence that DNA degradation in ancient bone is mainly driven by thermal age-dependent hydrolytic depurination driving rate-constant fragmentation over time. However, a separate analysis (24) found no significant link between sample age and the degree of fragmentation. Consistent with this latter finding, early ancient DNA research pointed to very rapid initial DNA decay followed by subsequent stabilization (25), rather than fragmentation as a rate-constant random decay process. Additionally, controlled experiments using qPCR with recently deceased tissues demonstrate a precipitous immediate decline of endogenous DNA content and/or quality, followed by stabilization hypothesized to be linked to the mineral environment of bone (26). This model likewise contradicts the idea that DNA decay can be thought of strictly in terms of exponential breakdown under a decay constant. In total, evidence has been presented for both a rate-constant decay model and a more age-independent scenario. Here, our meta-analysis points to the statistical decoupling of age and fragmentation. We aimed to validate this finding with three strategies:

First, we recognize that numerous sources of variance cannot be controlled in our meta-analysis across several studies—including sample excavation and storage conditions, wet lab and computational methods, and species and tissue types—and that these sources of variance have the potential to obscure subtle relationships. If major sources of inter-study confounding variance in DNA fragmentation were present, the result would likely be the dampening of any statistical relationship between natural variables and fragmentation as the *in situ* signal for fragmentation is lost. If age was a significant predictor of DNA decay along with thermal fluctuation and humidity, it is difficult to imagine that only the age relationship would be lost due to post-excavation handling and inter-study variation. Therefore, we suggest that confounding variance is not a parsimonious explanation for the lack of a clear age-fragmentation relationship in the presence of a robust environmental association. However, to test this possibility more directly, we restricted our analysis from 185 datasets across 21 studies to 97 Bronze Age human genomes generated from a single study (27). We thereby control for species, tissue type, and biases in sample preparation, and we consider a narrower timeframe and more constrained set of preservational conditions, eliminating several potential sources of confounding variance. Under the same linear model as above (Supplemental Methods), we find that exactly as in the broader dataset, thermal fluctuation and precipitation were strong predictors of fragmentation (respectively, *p* = 0.014 and *p* = 4.2 × 10^−4^; multiple r^2^ = 0.25), but age was still not a significant predictor of overall fragmentation (*p* = 0.420).

Second, we tested the fundamental assumption that from a single archaeological or paleontological site, DNA from older samples is expected to be more fragmented than from younger ones. While we initially analyzed data from 94 different sites, the meta-dataset includes 114 pairs of samples from the same site separated by at least 100 years. Thus for these 114 pairs where we can eliminate inter-site variation, the older sample is predicted to be the more fragmented sample a significant majority of the time under the fundamental assumption that fragmentation increases with age in a single environment. Given 114 pairs of samples, only 55 (0.48) satisfy this assumption (‘successes’). The null hypothesis of 47–67 successes (*p* = 0.05 calculated using the beta distribution) cannot be rejected, and indeed fewer than half of cases satisfy the basic assumption. By increasing the minimum age difference to 1000 years, we retain 55 valid pairwise comparisons and still observe no relationship between age and fragmentation, with only 27 (0.49) satisfying the basic assumption (null hypothesis at *p* = 0.05: 23–32 successes). We validated this approach by replicating the procedure with deamination, a known age-linked variable (24, and above). With deamination, we reject the null hypothesis and find a significant age effect as expected (131 comparisons possible, 80 successes (0.61); null hypothesis at *p* = 0.05: 55–76 successes).

Finally, we routinely observe complete deamination of all exposed cytosine residues. This saturation of measurable deamination has been described in several samples previously (23), and is observed in 14 out of the 185 (7.6%) datasets analyzed here, spanning 2kya to 500kya (Figure 4). However, complete deamination in single-stranded overhangs is incongruent with a rate-constant fragmentation model: If fragmentation followed a simple rate-constant process that would yield a robust association between thermal age and fragmentation, new overhangs would continually be exposed with the expectation of intact cytosine, suppressing the proportion of deaminated residues and preempting complete deamination. Even by simulating deamination rates tenfold faster than the most extreme of those estimated in our meta-analysis, deamination fails to converge to saturation under a rate-constant fragmentation model (Supplemental Methods). In total, observing complete deamination under a rate-constant fragmentation model would require that the deamination rate exceeds the fragmentation rate so that new overhangs are rapidly saturated with deamination—all exposed cytosine residues are rapidly converted to uracil. Under such extreme deamination rates, however, it is implausible that deamination would show such a robust correlation with age across samples as observed here and elsewhere (24).

**Figure 4.**
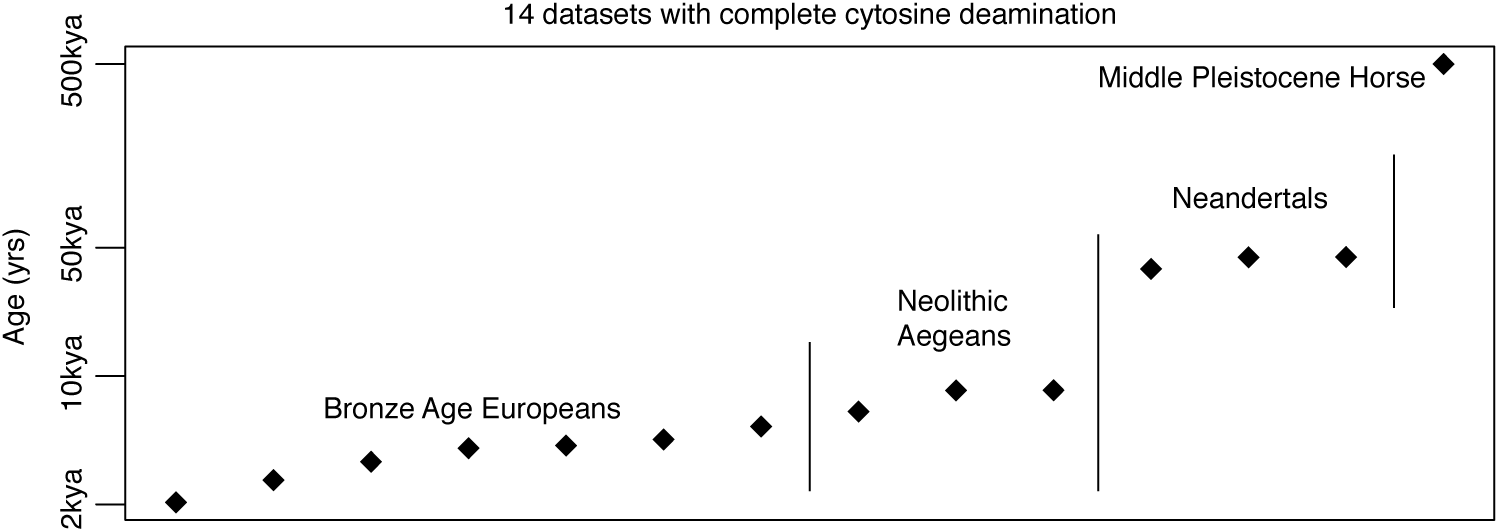
Fourteen samples included in the meta-analysis with saturated cytosine-to-uracil deamination in single-stranded overhangs.

We find strong validation that age does not predict DNA fragmentation in our meta-dataset. However, we recognize that DNA breakdown by hydrolytic depurination is a well-characterized and immutable chemical mechanism by which DNA decays exponentially according to first-order kinetics, producing a measurable half-life signal of molecular depletion (28). The mismatch between this predicted behavior and our findings indicates that the preservation state of ancient DNA is determined by multiple processes, and cannot be attributed to a simple fragmentation rate as suggested in a rate-constant fragmentation model. Instead, we propose a multi-stage DNA fragmentation model: First, physical and biotic stressors cause rapid breakdown of nucleic acids shortly after organism death. While microbes and cellular processes (e.g. autolysis and nuclease activity) rapidly degrade a large fraction of endogenous DNA—depending on tissue type and depositional environment—fragmentation appears to reach an initial threshold and then stabilize somewhat in contexts where DNA has the potential for long-term preservation.

The strong association of humidity and thermal fluctuation with DNA fragmentation suggests that processes like the loss of bioapatite surface area caused by diagenetic recrystallization and physical shearing effects of hydraulic fluctuations in bone, for example, may play a role in the initial breakdown process. Further, DNA may reach a size in bony contexts—the majority of our re-analyzed datasets—where it can penetrate the protective internal porosity of bone and gain some additional protection from the mineral environment. The counterintuitive result that DNA is sometimes better preserved in cooked than uncooked medieval bone may offer support for this scenario (29, although see 30 for further analysis of cremated bone). In our analysis, 15 plant samples from herbaria (31) fit with the overall fragmentation model—comparing fragmentation linear model residuals reveals no significant difference between plant samples and non-plant samples (Welch’s t-test, *p* = 0.44). However, they make up a very small fraction of the variation here, and because of the possible role of the mineral makeup of bone in DNA preservation across samples (26), we suggest that re-analysis of plant data across a much greater age range will be important in understanding any possible differences in preservation between plant and animal tissues. Over a short timespan, age-dependency in fragmentation has been documented in plant tissues (32), but the currently paucity of paleogenomic plant data currently precludes a comprehensive analysis spanning thousands of years. In total, our meta-analysis and model are necessarily focused on mammalian hard tissue (n=169 out of 185 datasets) given dataset availability. As more datasets are generated from diverse systems and tissue types, we expect further refinement of these general findings to reflect a more nuanced understanding behind the specific drivers of DNA diagenesis and factors underlying preservation. For example, DNA is integrated into hair during programmed cell death and keratinization leading to some amount of immediate shearing which might affect downstream processes (33). Thus ancient DNA in hair might warrant a modified set of expectations for preservation relative to bony tissue given a certain background environment. Recent experimentation comparing tooth cementum and petrous bone DNA diagenesis reinforces the necessity of integrating sample type information in assessing DNA degradation in the future (30).

We also find that in addition to the humidity and thermal fluctuation pattern, the degree of DNA fragmentation correlates strongly with base compositional biases. Specifically, datasets dominated by short fragments are significantly depleted of weakly-bonded nucleotide motifs (*p* = 6.79 × 10^−12^, r^2^ = 0.253; Figure 2; Supplemental Methods), suggesting that DNA breakdown follows predictable patterns with regard to microenvironment and nucleic acid biochemistry. Relatedly, we detected a histone-associated fragmentation bias (20) in the majority of our samples (n=112; Supplemental Methods), and we find that annual mean temperature is associated with the intensity of this pattern (*p* = 1.2 × 10^−5^, r^2^ = 0.16; Figure 2). Specifically, DNA breakdown in colder environments appears to more faithfully reflect cellular architecture and the *in vivo* genome context, whereas breakdown in warmer conditions is much less discriminant.

Previous research identified a strong age dependency in DNA recovery—assayed by quantitative PCR—in a controlled time-series of bone samples from a regional set of depositional sequences, and interpreted the result as evidence for an exponential decay process due to time-dependent DNA fragmentation (18). However, bulk diffusion of DNA—rather than rate-constant fragmentation—provides an equally parsimonious scenario for the observed qPCR signal. Specifically, the previous study estimated a 521-year half-life for a target fragment of 242bp in the tested environment (18). We estimate, however, that the same qPCR signal is consistent with bulk loss of 0.0013 of all remaining molecules per year as an alternative to rate-constant fragmentation (Supplemental Methods). As such, our results do not conflict with the previous experiment identifying a time-dependent decay behavior in relative copy number of a given fragment size. However, we propose that bulk DNA loss is congruent with both this qPCR signal and our meta-analysis, whereas exponential decay by fragmentation is not supported as the primary mechanism of DNA loss in our analysis. Therefore, we propose that much of the time-dependent nature of ancient DNA recovery may be due to bulk loss of DNA from tissue. Recent research focusing on the dense, non-vascularized petrous part of the temporal bone as a source of high endogenous DNA content (30, 34) demonstrates that targeting “semi-closed” systems with little opportunity for chemical exchange may be the best strategy to continue pushing the boundaries of DNA preservation by combating this diffusion process. This idea has also been robustly illustrated in studies dealing with DNA preserved in hair, which is thought to confer a protective micro-environment that impedes biological degradation, leaching, and possibly hydrolytic damage, and therefore often constitutes a good source of relatively high-quality endogenous DNA (33, 35).

We suggest that rate-constant fragmentation through hydrolytic depurination is seldom the limiting factor to long-term DNA preservation, but we offer some caveats: Fragmentation through depurination is a well-characterized process (28), and we do not propose that it is irrelevant for long-term DNA degradation. We suggest, rather, that the rate of this process is significantly slower than previously estimated in many ancient tissues (18), and the signal over the timespan re-analyzed here is overprinted by other factors in a multi-faceted breakdown process. Thus when estimating the value of ‘lambda’ for a dataset—the parameter describing fragment length distribution (Supplemental Methods)—we are analyzing the outcome of multiple processes rather than inferring a simple decay rate. Further, importantly, any paleogenomic meta-analysis is fundamentally limited to those scenarios in which DNA actually survives over Quaternary timescales, and so hydrolytic fragmentation as previously described might be a central mechanism for the total postmortem depletion of DNA in many tissues and conditions. That is to say, we can only analyze DNA that has survived, which may represent an abnormal mode of diagenesis. Our model for ancient DNA decay therefore necessarily speaks only to the special case in which conditions exist for long-term DNA survival. The immutable depurination process likely still imposes practical limits on DNA recovery in deep time, and recovering Mesozoic DNA, for example, remains extremely unlikely. However, semi-closed chemical exchange systems like the petrous bone, though rare, offer excellent potential for the long-term retention of DNA in tissues, and extraordinary preservational micro-environments created by chemical interactions have proven valuable for deep-time protein preservation (36). Breaching the current Middle Pleistocene age boundary of genomics seems entirely plausible.

## Acknowledgments

Research was supported by NERC Independent Research Fellowship NE/L012030/1 (to LK). We thank Ludovic Orlando, Beth Shapiro, and Kes Schroer for comments on an early version of the manuscript.

## Data availability

Analyses were based solely on publicly available datasets. Summary data are available for re-analysis as Supplemental Dataset S1. A tar file containing complete metadata and results from analyses, custom scripts, and run logs has been uploaded as Supplemental Dataset S2.

## Supplemental Methods

### Datasets and initial processing

We obtained unmapped (fastq) or mapped (bam) sequence reads from each of 185 publicly available ancient DNA datasets generated by shotgun sequencing without uracil removal, comprising anatomically modern humans (n=156; (27, 34, 35, 37–46)), herbarium plant samples (n=15; aligned to the host plant rather than the pathogen examined in ref (31)), Colombian and woolly mammoths (n=4; (7, 47)), neandertals (n=3; (48)), horses (n=5; (13, 49, 50)), and polar bear (n=1; (51)). We avoided data generated through target capture experiments to avoid possible hybridization biases introduced by misincorporated residues or read length variation. For unmapped samples, we used Flexbar (52) to trim adapter sequences in single-end read data, and PEAR (53) to perform adapter trimming and read merging in paired-end datasets. We used the bwa-backtrack algorithm within the Burrows-Wheeler Aligner (54) to map read data to the relevant reference genome, and collapsed duplicates using the rmdup function in SAMtools (55). We filtered all bam files for a minimum mapping quality of 20 using the SAMtools “view” function, and filtered for minimum read length of 20 using Unix tools. We separated nuclear and organellar reads (mtDNA in mammals, plastid DNA in plants) into separate bam files. For the mammoth samples mapped to the African elephant genome (n=4), we removed mitochondrial reads from the bam file and re-mapped the complete raw datasets to a woolly mammoth mitochondrial sequence (NC_007596.2). Following initial curation, we used mapDamage 2.0 (23) to estimate deamination and fragment overhang length distribution. We then estimated fragment length distribution, histone periodicity, and *k*-mer compositional biases using methods described below.

Sample latitude and longitude were used to estimate annual mean, minimum, and maximum temperature estimates, plus annual precipitation for each of the samples. These were taken from the WorldClim (56) current condition database using the R ‘raster’ package (57) at a resolution of 2.5 arc-minutes. In cases where specific site location was not available at the longitude/latitude level, Google Earth (accessed Nov 2015) was used to estimate longitude and latitude from details or site maps provided in the relevant publications. Location details and temperature estimates are given in Dataset S1. Climate estimates reflect modern climate conditions rather than a complete climate legacy over the timespan of each sample.

### Deamination estimation

We used mapDamage 2.0 (23) to estimate deamination in single-stranded overhangs, δ_s_, invoking default settings with the following exceptions: We subsampled large bam files to correspond with a 1 Gigabyte input file (∼10-20 million reads with typical dataset complexity and a human genome) using the mapDamage “-n” option. We analyzed the MCMC output from each sample using the ‘coda’ R package (58) to estimate an effective sample size (ESS) for each of the six variables estimated by the mapDamage simulation. ESS values are reported in supplemental Dataset S1. We enforced a minimum ESS of 200 in all variables to ensure MCMC simulation convergence, excluding nine datasets for deamination analysis. For libraries with highly asymmetrical 3’ and 5’ C-to-T mismatch observed visually in misincorporation plots, indicating the likely use of a non-proofreading DNA polymerase for library amplification—incapable of recovering uracils in template DNA—we re-ran mapDamage with the “––reverse” option to estimate damage from the 3’ end only. We noted extremely high deamination and overhang termination (*λ* in mapDamage) values in the output from Mammoth M4 (7), which suggested a much higher rate of deamination than even much older permafrost samples. However, that library is dominated by very short fragments ((7); summarized in the fragment *λ* plot in Supplemental Dataset S2), which we hypothesized could influence the mapDamage MCMC to over-estimate both parameters. We re-analyzed that sample considering only reads ≥40nt, yielding the damage parameter values reported in Dataset S1. The Saqqaq data (35) were mostly generated using a non-proofreading enzyme, but a small proportion of read files were reported to have been generated using a proofreading Platinum High Fidelity *Taq* polymerase (20). We mapped all Saqqaq read files from the Sequence Read Archive (n=218) to a human mitochondrial genome (EU256375.1), used PMDtools (59) to rapidly generate misincorporation plots, and visually inspected each for elevated 5’ C-to-T mismatch. This approach yielded two libraries apparently produced using a proofreading enzyme, one of which (SRR030983) was carried through for analysis. All mapDamage output files (run logs, plots, MCMC trace files, and summary statistics) from the 185 final runs are available in Supplemental Dataset S2. We then summarized a deamination rate for each sample according to the equation:

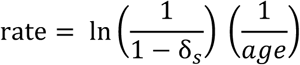

### Fragment length estimation (*λ*)

Fragment lengths are expected to form an exponential distribution under random breakdown. The distribution of DNA fragment sizes can therefore be summarized as *λ* (60), the single parameter of the exponential distribution. To estimate *λ*, we first summarized a frequency distribution table of fragment lengths. If a frequency spike was observed at the maximum fragment length—indicating fragments greater than the read length and an artifactual peak among reads with no adapter trimming—we re-estimated the maximum reliable fragment length as follows: Beginning with the longest fragment, we pruned the table back to the point at which the next shorter fragment was observed more frequently, eliminating up to 6 length values (mean = 3). We iterated over all ranges of at least 20 consecutive length values in the table, attempting to fit an exponential formula using the R function: *nls(y ∼ N*exp(-k*x))*, with starting values of *k*=0.05, *N*=0.1, and *λ* represented by the inferred value of *k*. We retained the top 5% of best fits on the basis of *p*-values obtained by summarizing the formula output in R, and estimated the value of *λ* from the table segment producing the best overall fit. We visualized the results in the top 5% of best fits to confirm a reasonable *λ* estimate (e.g. Figure S1). We observed fragment length heterogeneity in some cases, likely created by mapping biases, and occasional anomalous spikes in length frequencies that disrupted automated estimation of *λ*. Therefore, during visual inspection, we sometimes opted to override the inferred *λ* value by i) manually defining a range of fragment lengths over which to re-calculate *λ*, and/or ii) clipping artifactual frequency spikes by imposing a single frequency threshold value (e.g. Figure S1). All summary statistics and plots for *λ* estimation are available Supplemental Dataset S2, including run logs detailing manual override decisions. Perl and R code for lambda estimation are available Supplemental Dataset S2. During statistical comparisons, we analyzed the natural log of *λ*.

**Figure S1.**
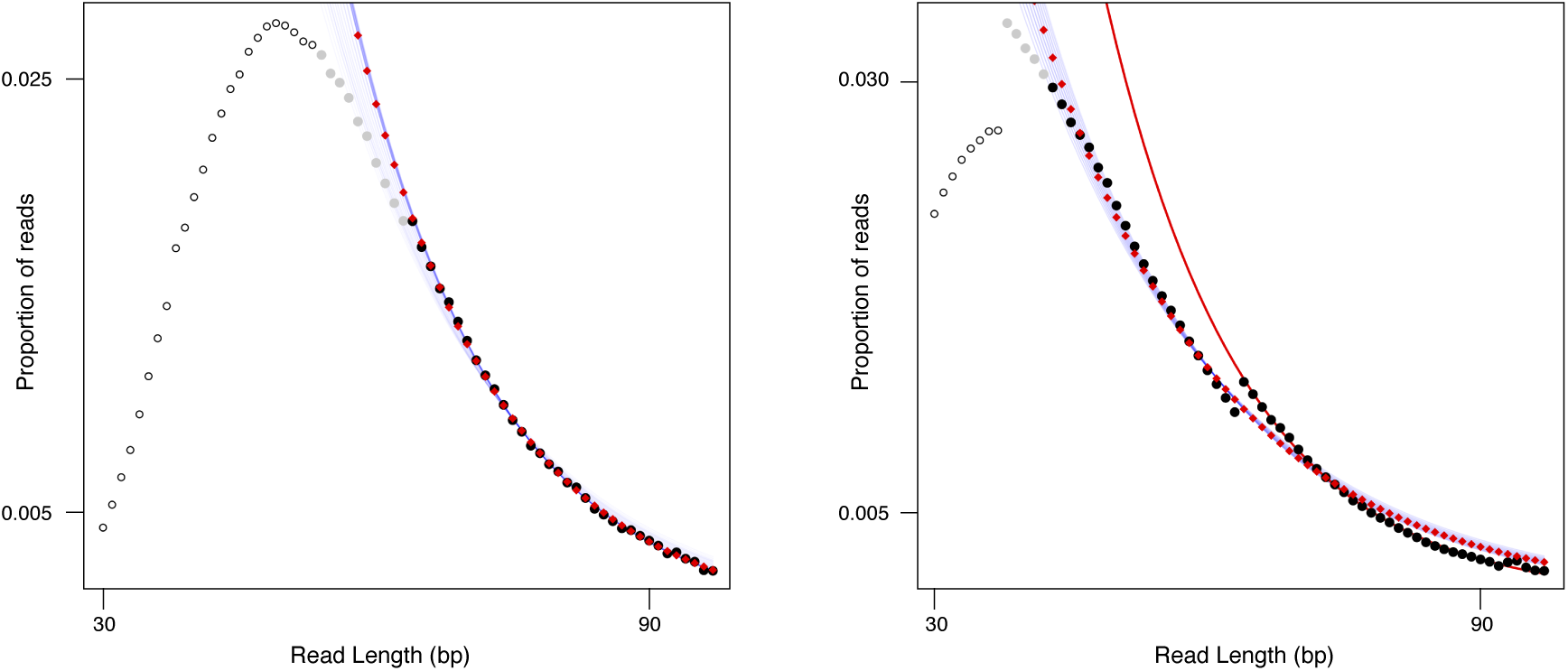
Automated estimation of *λ*. Each consecutive range of 20 length values is tested for a significant fit to the exponential distribution, the top 5% of best fits on the basis of p-value are retained, and the top hit is used to determine the value of *λ*. Left: gray shading shows all points in the top 5% of trials, and black points were used in the final estimation (best hit). Faint blue lines show the inferred exponential distribution under the *λ* values of the top 5% of trials, and red points show extrapolated frequency estimates under the best fit *λ* value. Right: In cases of length heterogeneity or other artifacts, automated estimation can be manually overridden by constraining the length range considered. In this case, *λ* is estimated using the range 65 to 90, and the red line shows the new estimate. Code to estimate lambda and generate these plots is available in Supplemental Dataset S2, as well as run logs and plots for 185 datasets analyzed here.

### Histone periodicity estimation

To estimate the intensity of a preserved histone signal, we analyze periodic deviations from a medium-range smoothing algorithm imposed on the fragment length frequency tables (Figure S2). Using a fragment length frequency table, we again eliminated the artifactual peak at the maximum length as above for *λ*, and trimmed the distribution to the innermost length values each representing at least 0.002 of the total fragments, as lower proportions were found to be noisy. Additionally, for artifactual spikes within the frequency table, we adjusted any single frequency greater than 1.5x the midpoint of its flanking neighbors down to the midpoint. This approach affected only artifacts, and not the underlying distribution. We then fit a locally weighted scatterplot smoothing (lowess) curve with the R ‘lowess’ function, using a smoothing span of 20 to normalize over histone periods expected to be ∼10nt (20). We then removed 4 additional length values from each end of the distribution to eliminate increased terminal deviations from the lowess curve.

We observed that in samples with a histone signal, the deviation of the observed values from the lowess curve is best approximated by a series of local exponential functions with the midpoint of a complete histone period set to *x*=0 (Figure S2). That is, frequency values across a single histone period form a parabolic curve when normalized for overall fragmentation described by *λ*. Therefore, we tested for this pattern in all subranges of 8–12 consecutive length values in the table, setting the midpoint of each subrange to *x*=0 and using the observed value divided by the lowess values for the y axis. We used the R function 
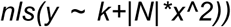
, with starting values of *k*=1 and *N*=0.1, so that *k* should deviate minimally from 1 to absorb noise, and positive values of *N* provide a metric of signal intensity. That is, *N* increases linearly with the degree of observed frequency deviation from the lowess curve at local maxima. We retained the starting position, range lengths, and *N* values of all significant fits on the basis of *p-*value. We then used an optimized one-dimensional *k*-means clustering algorithm in the R ‘Ckmeans.1d.dp’ package (61) to localize strongly significant starting locations of histone periods. For all adjacent (*i, j*) pairs of cluster positions representing the putative best locations for histone peaks, a periodicity coefficient was calculated: If *j* – *i* ≥ 8 and *j* – *i* ≤ 12, the coefficient increases by 1/(*n* clusters -1). Otherwise, if some value *v* in 2 through (range of values)/10 satisfies (*j* − *i*)/v ≥ 8 ⋃ (*j* − *i*)/v ≤ 12, coefficient increases by (1/v)/(*n* cluster − 1). Thus, given a minimum of three clusters and a minimum periodicity coefficient of 0.3, cluster position values satisfying the requirement could include 10,20,40 (coefficient = 0.75); 10,20,50 (0.67), 10,30,50 (0.5), or 10,40,70 (0.33), but not 10,50,80 (0.29).

**Figure S2.**
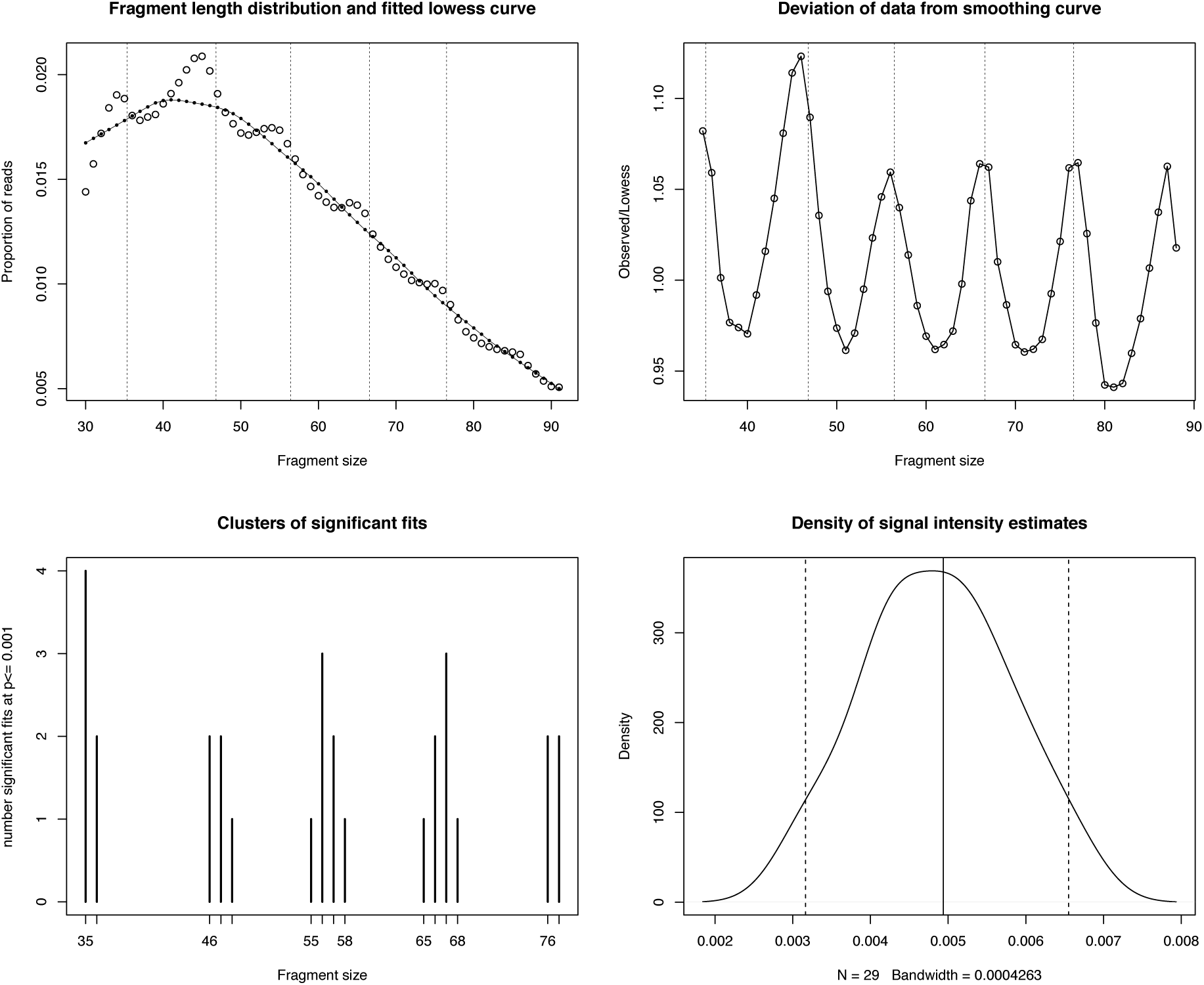
Histone periodicity estimation pipeline. To estimate the intensity of the histone periodicity signal, a lowess curve is fit to the fragment length distribution (top left), and the deviation of the data from the smoothing curve is calculated (top right). The distribution is then scanned for significant local parabolic regions, and significant starting locations are identified using a 1-dimensional clustering algorithm (bottom left). Finally, the density distribution of the intensity variable is summarized, and the median is recorded as an estimate of histone signal intensity. Code for histone intensity estimation is available in Supplemental Dataset S2, along with run logs and plots for 112 successfully estimated samples.

To optimize histone signal estimation, we used the organellar datasets (which lack histones *in vivo*) to calibrate model parameters against false positive results (Figure S3). We permuted the minimum number of significant fit clusters detected (2, 3, 4), a minimum observed proportion of all plausible histone periods given the range of values analyzed (0, 0.1, 0.2, 0.4), the minimum periodicity coefficient (0.2, 0.3, 0.4, 0.5), and a minimum *p*-value threshold for significant exponential fit (0.05, 0.01, 0.001). We summarized the number of nuclear and organellar datasets satisfying the requirements of 144 separate model permutations (*n* = 53,280 total iterations). p-value, minimum cluster count, and minimum periodicity coefficient proved the best predictors of false positive rates, accounting for 79% of the variance in false positive rate under a simple linear model (Figure S3), while proportional number of clusters did not add predictive power. Under a range of parameter values, we were able to estimate nuclear histone signal intensity with no organellar false positives in up to 112 of the 185 samples for a given model. Using overly relaxed conditions, estimates could be obtained in 167 samples, but at the expense of specificity, with over half of the organellar datasets (n=101) yielding false positives. Given a 5% allowable false positive rate in the single model with the highest ratio of nuclear to organellar estimates, we were able to recover 138 nuclear estimates. We recorded the median value of the *N* intensity parameter for all samples under strict conditions with no organellar false positives (n=112) for analysis (Dataset S1 ‘Histone_Intensity’ for estimates; see Supplemental Dataset S2 for pdf and run files). Notably, the Thistle Creek horse genome (50) displays a clear short-range periodicity on visual inspection, but at about half the normal length (∼5bp). The reason for this behavior is unclear, but this distribution violates the model assumption of a ∼10bp period, and therefore this sample only ever presented as a likely false positive during model calibration.

**Figure S3.**
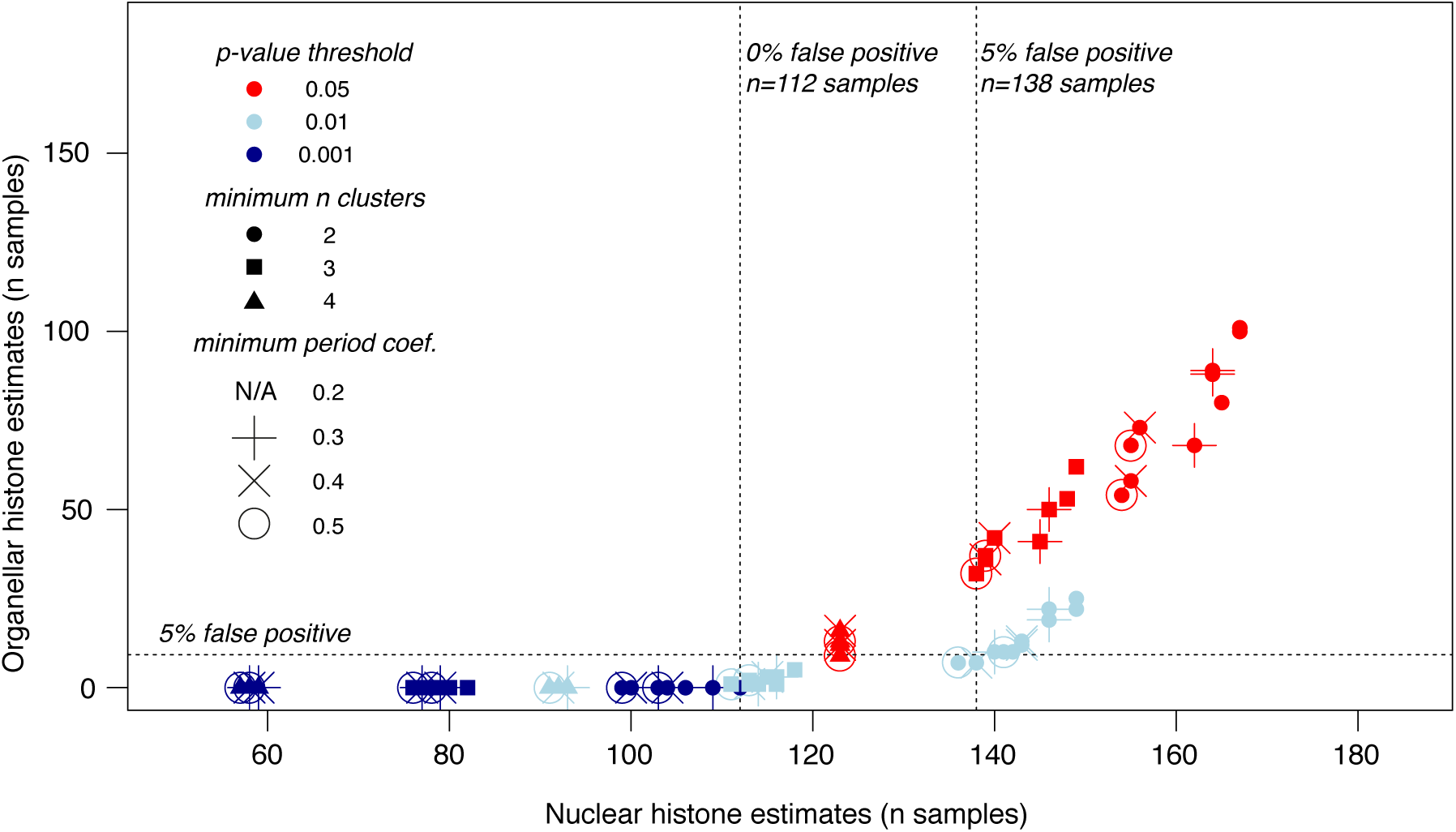
Plot shows the number of successful nuclear and organellar automated histone intensity estimates across 144 model configurations, where each point represents the same 185 samples under a different model. Organellar estimates are assumed to be false positives, so the y-axis provides a proxy of the false positive rate that increases with relaxation of parameters. Single models can be optimized to yield up to 112 nuclear estimates before organellar false positives are detected (n=64 models with no false positives), and the p-value of the exponential formula nonlinear least squares fit used to localize parabolic regions is the best determinant of false positive rate. By accepting a 5% false positive rate, we can recover estimates for 138 samples under the best single model.

### Base composition

We summarized 8-mer frequencies in each reference genome, excluding soft- and hard-masked repeat regions, using a custom perl script. We then summarized 8-mer frequencies in the bam files from sequence reads in mapped orientation to match the reference. For each 8-mer, a simple enrichment factor was calculated as (frequency in reads)/(genomic frequency). The enrichment and depletion of ancient DNA motifs is affected by a complex range of conditions, as suggested by clear multimodality in the distribution of 8-mer enrichment factors (Figure S4). Additionally, *in vitro* variables further bias the datasets through penalizing GC-extreme reads, for example (62), and chromatin modeling and nucleosome occupancy is expected to have differential effects on the protection and survival of coding vs. non-coding DNA. To isolate these effects, we calculated a simple GC proportion for each 8-mer, and based on known systematic sequence complexity biases among genomic element types (63), we calculated 8-mer Shannon entropy (*H*) using the following formula after ref (64):

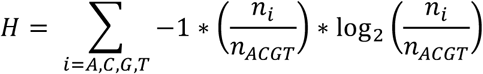

Finally, we calculated a simple kmer enthalpy as the sum of all dimer base-stacking energy values (kcal/mol/dimer) reported in table IX of ref (65), using the values from the “corrected optimized potential” method. We first visualized kmer enrichment in relation to all three sequence variables—GC content, entropy, and enthalpy—and noted extensive variation among samples as expected (e.g. Figure S4, kmer summary files for all datasets, code for 3d scatterplots, and code for enthalpy-biased kmer frequency estimation are available in Supplemental Dataset S2). For example, GC content and entropy are parabolically related by definition—equal base frequencies are required for maximum entropy. As such, disentangling enrichment of high-entropy kmers from *in vitro* penalization of GC-extreme kmers is intractable, and kmer enrichment patterns varied extensively in terms of entropy and GC-content. However, we noted a strong relationship between enrichment and enthalpy in several samples (Figure S4), and therefore we opted to isolate enthalpy from GC content and entropy for analysis as follows:

In total, there are 52 unique GC-content – Shannon entropy combinations for DNA 8-mers. Each of the 52 configurations of invariant GC and entropy values represents between 2 and 6720 kmers (*n* = 65,536 total kmers) representing a distribution of enthalpy values and enrichment factors. From within each of the 52 GC–entropy configuration bins, we reiteratively selected random pairs of kmers up to the total number of kmers in the bin (i.e. given a bin containing 32 kmers, 32 random pairs were selected, and a total of 65,536 pairs are drawn from the dataset). We then compared enrichment factors and enthalpy values between the two kmers. Given a “success” if the higher-enthalpy kmer was also the more enriched kmer, as hypothesized, we incremented a counter, and decremented the counter for failures. Following iterations, we recorded the value of the counter divided by the number of trials, and used this value as the test statistic to compare with environmental variables, where positive values indicate overall enrichment of higher-enthalpy motifs (Dataset S1, “Enthalpy_bias”).

**Figure S4.**
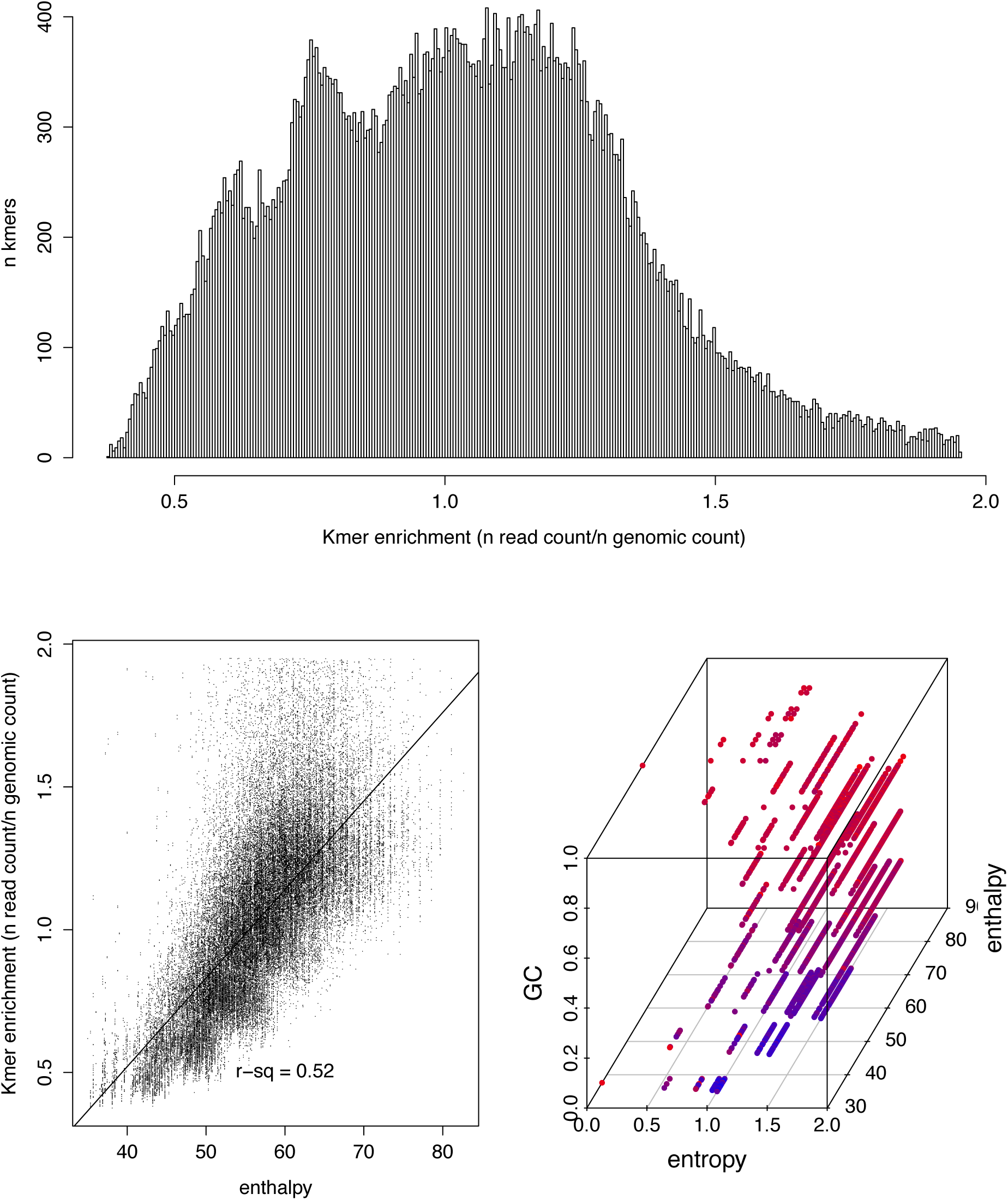
Kmer enrichment in the ∼13kya Anzick (clovis) genome (40). Top 5% of enrichment values were excluded prior to visualization (none were excluded during analysis). Top: Histogram of kmer enrichment factors showing multimodality. Bottom left: strong correlation between kmer enthalpy and kmer enrichment. Bottom right: kmer enrichment compared with simple GC content, enthalpy, and Shannon entropy. Red points represent enriched kmers, blue points represent depleted kmers. Kmer enrichment files and code to estimate bias available in Supplemental Dataset S2.

### Linear Models

We carried out multiple linear regression analyses to test for relationships between preservation parameters (above) and environmental variables. Specifically, we tested in turn for significant predictors of each damage parameter using a linear model analysis with four independent variables: annual mean temperature, annual temperature fluctuation, annual mean precipitation, and the natural log of sample age using the R ‘lm’ function (66). Results presented are from analysis of the nuclear datasets after excluding the organellar data. We chose to work with δ_s_ directly instead of the first-position C->T misincorporation frequency often used to describe deamination (e.g. (24)), as the latter is a compound statistic reflecting δ_s_ and the overhang lambda statistic estimated using mapDamage.

### Endogenous DNA loss vs. DNA fragmentation

Decay behavior in the molarity of a target DNA fragment over time in a qPCR assay could be attributed to either rate-constant fragmentation (18) or bulk loss of endogenous DNA. In the example from ref (18), researchers inferred a nucleotide fragmentation rate (*k*) of 5.5 × 10^−6^ damage events per nucleotide per year on the basis of a strong relationship between age and 242bp target molarity across a large set of mass-standardized bone samples. Under a rate-constant fragmentation model where *k* = 5.5 × 10^−6^, the probability of retaining any fragment of length *L* per year is the probability that no random breakage event occurs at any of its sites, or (1-*k*)^*L*^. The probability of fragment loss (*k*_*L*_) is the opposite: 1-(1-*k*)^*L*^. For a 242bp target, therefore, *k*_242_ = 1.33 × 10^−3^—each year 0.0013 of remaining 242bp templates are severed on average. However, this model assumes no time-dependent loss in DNA by mechanisms other than fragmentation, but if fragmentation stabilizes and bulk depletion of endogenous DNA continues, a similar pattern would result. Specifically, if each fragment has a 0.0013 probability of being lost to bulk DNA movement rather than fragmentation, the same qPCR signal of decreasing target molarity over time would result. Given that our meta-analysis shows no association between sample age and directly-estimated *λ* values, we propose that loss of endogenous DNA is the more parsimonious mechanism of half-life behavior in 242bp molecule loss than rate-constant fragmentation.

### Simulating DNA fragmentation and deamination

We hypothesize that a time-dependent fragmentation process is incongruent with the observation of total cytosine deamination in single-stranded overhangs (δ_s_ = 1). We therefore carried out a set of simulations to show the effects of varying the fragmentation rate on the proportion of observed deaminated residues. Simulations were executed using a custom perl script. Beginning with a *λ* value of fragmentation (e.g. 0.013 to 0.157 range from our meta-analysis), we infer the number of random fragmentation events necessary to yield the lambda value as *λ* x (total length of all fragments). We then randomly select a simulated number of imposed total fragmentation events from a Poisson distribution, using the exact number of fragmentation events as the Poisson lambda parameter. We pre-allocate the selected number of breakage events—without replacement, as breakage is impossible twice at the same location—to locations in a population of starting molecules. Breakage occurs at zero-width sites between simulated residues in our simulation, such that molecules can be reduced to 1nt but not lost completely without additional parameters. We then allocate fragmentation events to timing bins by sampling from the probability density function of a beta distribution with the α parameter held at 1, and the β parameter varying ≥ 1 to introduce a changing rate profile: β = 1 describes a constant rate of fragmentation, and higher values of β describe increasingly skewed scenarios where fragmentation occurs in the early cycles of the simulation. For example, when α = 1 and β = 10, roughly 90% of fragmentation has occurred when 20% of time has passed. Alternatively, α = 1 and β = 1 describes a uniform distribution of fragmentation where the rate parameter *k* = *λ*/time, as per ref (18). To complete setup, we impose a δ_s_ value and calculate a deamination rate as described above in terms of probability of deamination per site per year, and impose a value to describe fragment overhangs per mapDamage 2.0 (0.3 in our simulation). Crucially, it is impossible to infer a deamination rate at δ_s_ = 1, as the equation would require taking the natural log of 0. However, we can impose extreme deamination rates separately if desired.

We then carry out a forward simulation through cycles of drawing from the randomized predetermined breakage sites according to the timing bin allocations, and introduce new single-stranded overhangs at newly broken sites by sampling randomly from a Poisson distribution described by the overhang *λ* value. Following each breakage cycle, each overhang is subjected to a round of deamination according to the rate calculated from δ_s_, where each site is given the opportunity to undergo deamination if a pseudo-random number (0 ≤ *x* ≤ 1) falls below the rate value. Finally, at the end of each deamination cycle, we summarize the current fragmentation *λ* value as 1/(mean current fragment length), and the current δ_s_ value as the (number of deamination sites)/(all overhang sites). When α = 1 and β = 1, *λ* increases linearly to approach the imposed *λ* value, while with higher β values, alternative patterns occur. Our simulations demonstrate that uniform time-dependent breakdown depresses the value of δ_s_ through constant introduction of unmodified single-stranded overhang, even at extreme estimates of deamination rate.

**Figure S5.**
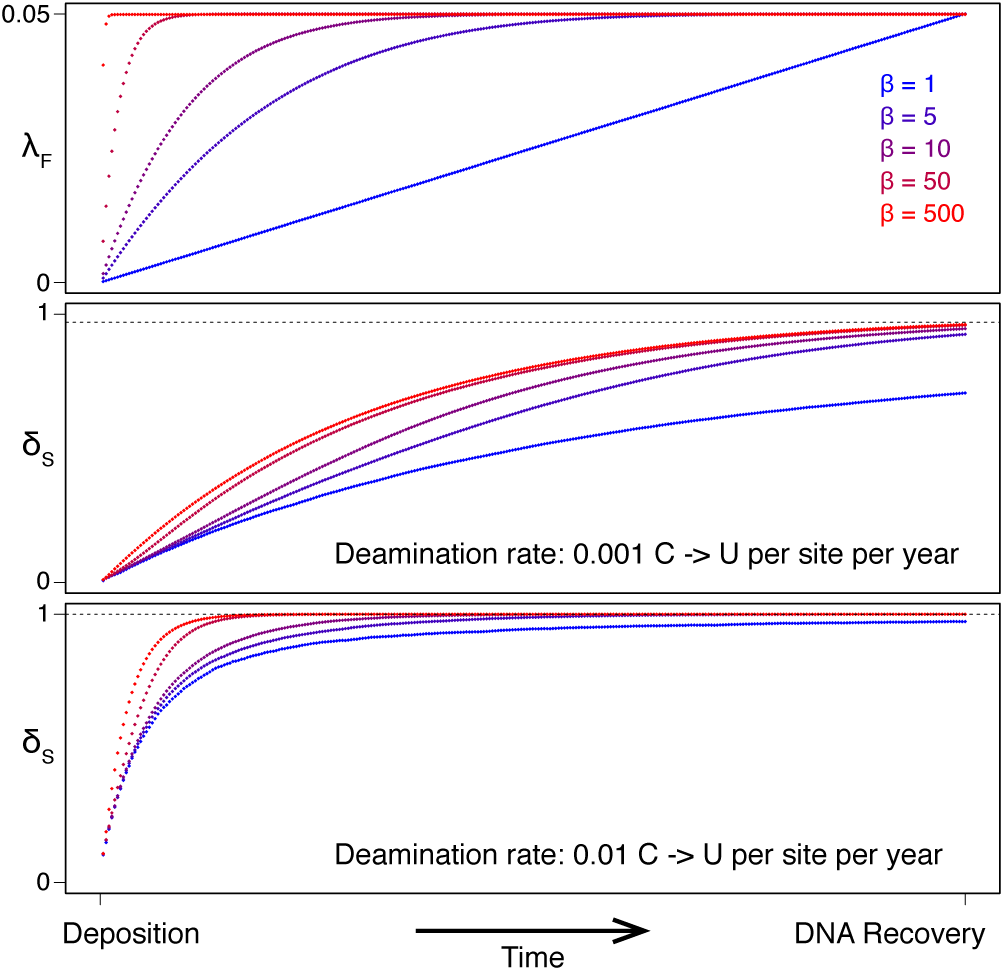
Simulation of deamination under variable fragmentation models. Top: Under a rate-constant model (β = 1), DNA breaks occur steadily over time and *λ*_F_ is assumed to increase linearly from the time of organism death to the time of DNA recovery. Increasing values of β impose a bias so that breakage occurs early and then stabilizes at the measured *λ*. For example, under β = 10, 90% of fragmentation occurs in the first ∼20% of time. Extreme values of β describe an instantaneous (entirely time-independent) fragmentation process. Middle: The effects of varying the breakdown rate distribution over time on the accumulation of deaminated residues and the saturation of δ_s_. The deamination rate of 0.001 C -> U per site per year was imposed based on the fastest empirical deamination rate observed from the 185 samples, and the dashed line marks the δ_s_ value of 0.97 associated with that sample. Bottom: By increasing the rate tenfold, expected saturation to δ_s_ = 1 (marked by the dashed line) is reached in all scenarios except under a rate-constant fragmentation model. This suggests that δ_s_ = 1 observed in paleogenomic datasets is incongruent with rate-constant DNA fragmentation.

### Deamination prediction from temperature

We calculated a density distribution of deamination rates using default parameters and bandwidth in the R ‘density’ function, and creating a weighting vector where each point’s weight value was calculated as 1-(local density/maximum density). We then fit a weighted linear regression between temperature and deamination rate using the R ‘lm’ function with the ‘weight’ option invoked using the above weighting vector. We used the R ‘predict’ function to predict a rate and 95% confidence intervals at temperature values of -20°C, 0°C, and 20°C. The R code to replicate the analysis and re-create Figure 3 from Dataset S1 is available in Supplemental Dataset S2 in the “expedient code” file.

